# Site-specificity of streptococci in the oral microbiome

**DOI:** 10.1101/2021.11.29.470446

**Authors:** Anthony R. McLean, Julian Torres-Morales, Gary G. Borisy, Jessica L. Mark Welch

## Abstract

Patterns of microbial distribution are determined by as-yet poorly understood rules governing where microbes can grow and thrive. Therefore, a detailed understanding of where bacteria localize is necessary to advance microbial ecology and microbiome-based therapeutics. The site-specialist hypothesis predicts that most microbes in the human oral cavity have a primary habitat within the mouth where they are most abundant. We asked whether this hypothesis accurately describes the distribution of the members of the genus *Streptococcus*, a clinically relevant taxon that dominates most oral sites. Prior analysis of 16S rRNA gene sequencing data indicated that some oral *Streptococcus* clades are site-specialists while others may be generalists. However, within complex microbial populations composed of numerous closely-related species and strains, such as the oral streptococci, genome-scale analysis is necessary to provide the resolution to discriminate closely related taxa with distinct functional roles. Here we assess whether individual species within this genus are generalists using publicly available genomic sequence data that provides species-level resolution. We chose a set of high-quality representative genomes for *Streptococcus* species from the human oral microbiome. Onto these genomes, we mapped short-read metagenomic sequences from supragingival plaque, tongue dorsum, and other sites in the oral cavity. We found that every reliably detectable *Streptococcus* species in the human oral cavity was a site-specialist and that even closely related species such as *S. mitis*, *S. oralis*, and *S. infantis* specialized in different sites. These findings indicate that closely related bacteria can have distinct habitat distributions in the absence of dispersal limitation and under similar environmental conditions and immune regimes. These three species also share substantially the same species-specific core genes indicating that neither taxonomy nor gene content are clear predictors of site-specialization. Site-specificity may instead be influenced by subtle characteristics such as nucleotide-level divergences within conserved genes.

## Introduction

Understanding the distribution patterns of bacteria across habitats is a fundamental building block of microbial ecology. In the human microbiome, the rules that determine whether a bacterium can colonize a given site are fundamental to understanding the role of the microbiome in health and disease and its potential as a therapeutic target. The Human Microbiome Project (HMP) was designed to establish a high-resolution baseline for similarities and differences in microbiome composition from individual to individual and site to site (Turnbaugh et al. 2007). Together with previous cultivation-based and cultivation-independent studies, the HMP demonstrated that bacteria occupy characteristic habitats: bacteria that are most abundant in the human gut tend to be rare on the skin or in the mouth, and vice versa (Costello et al. 2009, HMP Consortium 2012). Thus, most bacteria are found predominantly in one broad habitat type. Not yet clear, however, is what features of the habitat determine which bacteria can thrive, how finely subdivided the habitats are, and what range of micro-habitats each bacterium can occupy.

For addressing these questions, the human oral cavity provides a natural experiment with many replicates and built-in controls. The distinct surfaces in the mouth (including enamel as well as keratinized, non-keratinized, and specialized mucosa) represent distinct potential microbial habitats that are spatially adjacent, with minimal barriers to microbial dispersal (Proctor and Relman 2017, Mark Welch et al. 2020). Each human individual is an island whose mouth has undergone the process of colonization. The bacteria inhabiting the same mouth are exposed to the same host diet, behavior, and immune regime, controlling for many of the variables that might influence microbial community composition. The composition of the oral microbiome is relatively well-understood, with a curated database (Dewhirst et al. 2010) identifying ~700 bacterial species resident in the mouth. The majority of these species can be cultivated in the laboratory. The HMP (HMP Consortium 2012), as well as independent research efforts, has generated sequenced genomes for most of the cultivable oral microbes as well as shotgun metagenomic sequence data sampled from a variety of sites within the mouth for several hundred individuals. Thus, the knowledge base exists to support a systems-level study of the habitat distribution of the oral microbiota.

The site-specialist hypothesis for the oral microbiota was developed based on 16S rRNA gene sequence data from the HMP as well as prior cultivation and cultivation-independent studies (Gibbons et al. 1963, Gibbons et al. 1964 A, Gibbons et al. 1964 B, Gordon and Gibbons 1966, Gordon and Jong 1968, Frandsen et al. 1991, Mager et al. 2003, Aas et al. 2005, Zaura et al. 2009, Human Microbiome Project Consortium 2012, Huse et al. 2012, Peterson et al. 2013, Mark Welch 2014, Eren et al. 2014, Hall et al. 2017, Bernardi et al. 2020). These studies showed that whether a taxon appears to be a generalist or a specialist depends on the resolution of the analysis: at the genus level, most oral bacteria have representatives throughout the mouth, but at the species level they are site-specialists; most species preferentially colonize certain regions of the mouth (Mark Welch et al. 2019). This primary habitat for a species can sometimes be narrowly defined; for example, some bacteria are abundant only on the keratinized gingiva (Eren et al. 2014) and one, *Simonsiella mulleri*, appears to live exclusively on the hard palate (Aas et al. 2005, Eren et al. 2014). Often, however, the primary habitat is broader and consists of a group of sites; for example, many bacteria are specialists for both supra and subgingival dental plaque, others for the tongue dorsum, palatine tonsils, and throat (Eren et al. 2014, Mark Welch et al. 2019). These distribution patterns suggest specialized adaptation for a subset of sites within the mouth.

A major exception to this pattern is found in the genus *Streptococcus*, which contains both specialist and apparent generalist taxa. *Streptococcus* is the most abundant genus in the oral cavity (Dewhirst et al. 2010, Hsu et al. 2012, Mager et al. 2003). As primary colonizers, oral streptococci play important roles in biofilm formation (Jenkinson 1994, Li et al. 2004). Some members of the genus contribute to the progression of disease while others help maintain the health of their host (Abranches et al. 2018). Thus, the spatial distribution of this genus is a critical feature of oral ecology. Some species of oral streptococci are so closely related that short regions of the 16S rRNA gene alone cannot necessarily distinguish them. 16S sequences are notably insufficient for differentiating species within the Mitis group (Jensen et al. 2016, Croxen et al. 2018, Velsko et al. 2019). Consequently, many studies relying on 16S sequencing data only distinguish between a handful of oral *Streptococcus* operational taxonomic units (OTU) (Zaura et al. 2009, Huse et al. 2012, Eren et al. 2014, Hall et al. 2017). The analysis of Eren et al. (2014), distinguished 11 OTUs for around 30 known oral *Streptococcus* species and indicated that the *Streptococcus* genus includes both site-specialist taxa and an apparent generalist, the group containing the abundant oral commensal *S. mitis* and its close relatives *S. pneumoniae, S. oralis* and *S. infantis* (Mark Welch et al. 2019).

Here, we test the site-specialist hypothesis for each human oral *Streptococcus* species using whole-genome data combined with shotgun metagenomic sequence data. Our results show that each of these species demonstrates site-specificity within the mouth. These findings indicate that closely related bacteria can have distinct habitat distributions in the absence of dispersal limitation and under similar dietary and immune regimes. Further, distinct distributions occur despite whole-genome analysis showing no major differences in gene content or functional annotation, indicating that subtle differences in genomic sequence can have ecologically significant effects.

## Results

### Identification of representative genomes and species-level groups

Our strategy for assessing whether *Streptococcus* species are site-specialists or generalists was to obtain publicly available short-read metagenome sequencing data from samples from the mouth and align these samples to high-quality sequenced genomes of *Streptococcus* species using short-read mapping. We first constructed a set of genomes that were accurately identified to species and distributed as evenly as possible across sequence space. From genomes of oral streptococci in the NCBI Reference Sequence Database (RefSeq), we selected a set in which each genome shared no more than 95% ANI with any other genome (see “Methods” for details.) For some species, all sequenced genomes available at NCBI shared an average nucleotide identity > 95%; these species were each represented by a single genome in our set (e.g., *S. mutans, S. pyogenes, S. agalactiae*, and *S. salivarius* in Fig. S1). Other species were represented by multiple genomes. As has been previously reported, many genomes deposited into RefSeq for the Mitis group streptococci have questionable species designations, due to factors including high intra-species diversity relative to inter-species diversity as well as frequent horizontal gene transfer and recombination events between species (Chi et al. 2007, Donati et al. 2010, Jensen et al. 2016, Croxen et al. 2018, Velsko et al. 2019). Therefore, we checked the species identifications of the reference genomes by constructing a phylogenomic tree based on the concatenated amino acid sequences of 205 single-copy core genes present in all the genomes and by evaluating the ANI between all genomes. Numerous Mitis group genomes clustered within a species different than their NCBI designation (Fig. S1). We assigned these genomes new species designations reflecting their clade in the phylogenomic tree to obtain a usable set of reference genomes for metapangenomics and pangenomics.

Genomes of different species segregated into distinct groups rather than representing a continuum of relatedness, even among the closely related *S. mitis, S. oralis*, and *S. infantis*. The grouping of genomes according to phylogenomics was consistent both with relatedness as indicated by the ANI values and with prior phylogenies constructed with genomes identified as Mitis group species in NCBI (Fig. S1, Fig. 1). The genomes of most species formed monophyletic clades sharing 90-95% ANI. Exceptions included the *S. pneumoniae* and *S. pseudopneumoniae* type strain sequences, which fell within the *S. mitis* clade, and the *S. peroris* type strain sequence which was placed within the *S. infantis* clade consistent with phylogenies constructed for members of the Mitis group (Chi et al. 2007, Jensen et al. 2016, Kilian and Tettelin 2019). The combination of ANI and phylogenomics was insufficient to distinguish *S. pneumoniae* and *S. pseudopneumoniae* genomes from *S. mitis* because *S. pneumoniae* and *S. pseudopneumoniae* are effectively sub-clades within *S. mitis* (Jensen et al. 2016, Croxen et al. 2018, Velsko et al. 2019) and both species share > 93% ANI with some *S. mitis* strains (Fig. 1) (Croxen et. al 2018). To identify *S. pneumoniae* and *S. pseudopneumoniae* we aligned species-specific marker sequences for *S. pneumoniae* and *S. pseudopneumoniae* (Croxen et al. 2018) to all the reference genomes. This alignment resulted in the identification of a single genome representing *S. pneumoniae* and a single genome representing *S. pseudopneumoniae*, the type strain in each case.

**Figure 1:**
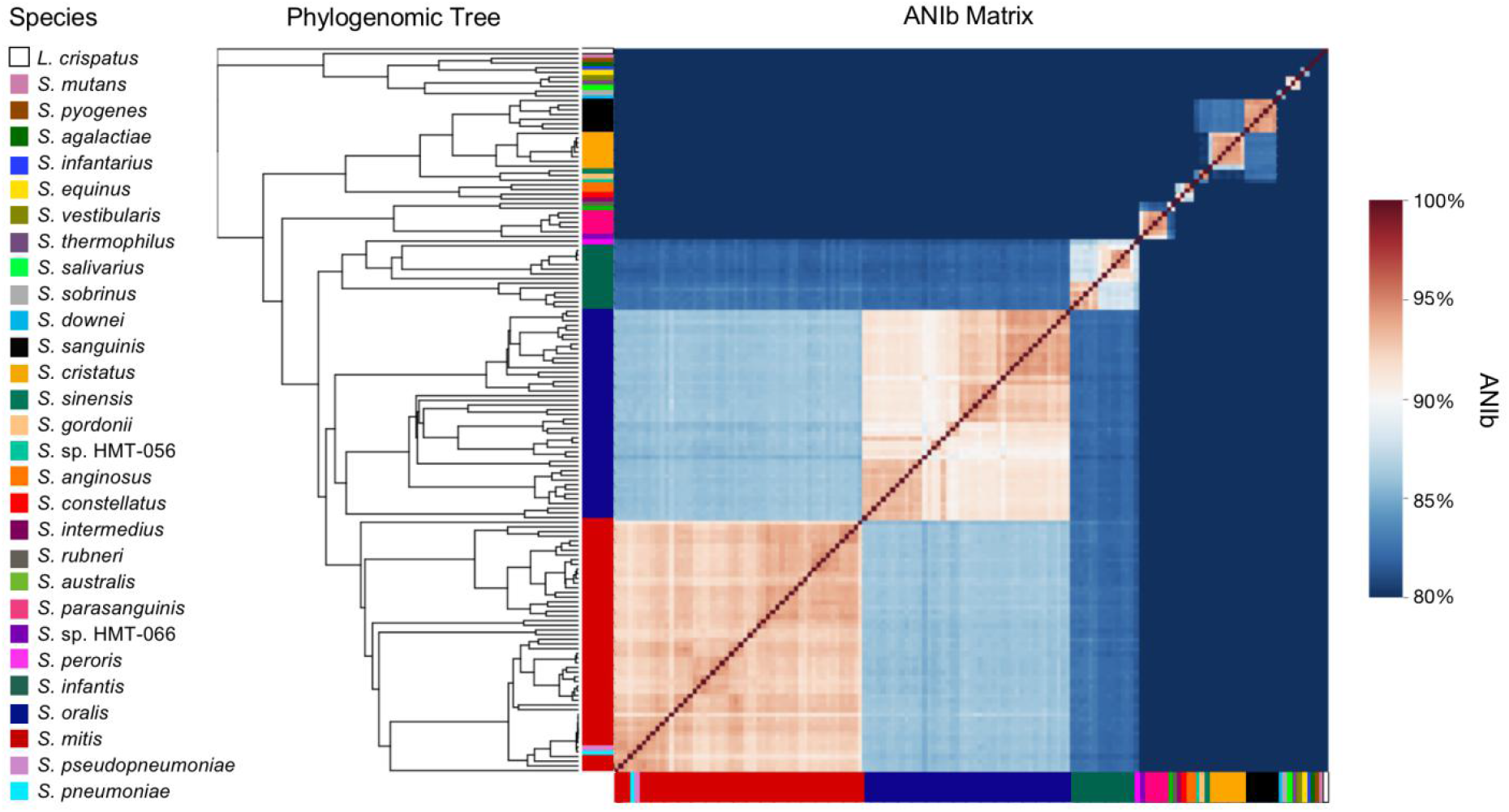
Clustering of genomes within the phylogenomic tree based on single-copy genes is consistent with relatedness based on average nucleotide identity. The phylogenomic tree was constructed using 205 single-copy genes core to the oral streptococci. The matrix displays the ANI calculated with the BLAST method (ANIb) between each genome in the tree and every other genome in the tree. The genomes are color-coded by species and arranged according to their placement in the phylogenomic tree.

### Tests using computationally generated short reads indicate that cross-mapping occurs at low levels and generally to closely related species

The accuracy with which a short-read mapping strategy can link a metagenomic sequence with its source species is limited by the degree to which sequences in different target species are an equally good match to the metagenomic read. In the mapping process, reference genomes from isolated strains of *Streptococcus* spp. act as bait to attract reads from the complex mixed population found in the mouth. The number of reads mapped should permit an accurate estimate of species composition if the short reads from one strain map much more frequently to the genome of another strain of that same species than to a genome from a different species. To test the accuracy of this expectation, we generated simulated short-read samples and mapped them to the selected set of *Streptococcus* spp. reference genomes. To generate each simulated read set, we computationally generated short reads from a single reference genome so that the simulated short reads covered the template genome to a mean depth of 100x across all nucleotide positions. As templates, we chose type strain genomes that were already in the reference set of genomes to which the reads were mapped; all short reads from these genomes can find an identical match to their source genome in the reference set but some may map instead to identical regions in other genomes. Additionally, we chose as templates genomes that were not present in the reference set but were from the same species and shared at least 95% ANI with a genome in the reference set, a situation that more closely approximates the expected composition of natural samples from the mouth. As non-*Streptococcus* spp. controls, we also used type strain genomes from other major human oral genera. We mapped the simulated samples to the reference genome set competitively (i.e., each read was compared against a file containing all the reference genomes, so that nucleotide positions across all genomes competed to match the read). In cases where a species was represented by more than one genome, we calculated the sum of the mean coverage of all the reference genomes belonging to that species and report this species-level coverage value.

Results indicated that mapping accurately identified the source species of reads. When the simulated sample used a template that was itself in the reference genome set, in most cases 98-99% of the reads mapped to the correct species (Fig. S2 and Table S2). Most cross-mapping was to closely related species, e.g., between *S. salivarius* and *S. vestibularis*, or between *S. intermedius, S. constellatus*, and *S. anginosus*. When the simulated sample used an oral *Streptococcus* spp. template not in the reference genome set, some reads failed to map to any genome in the set, but among the reads that did map, more than 80% mapped to the correct species except for *S. australis, S. constellatus, S. pneumoniae*, and *S. pseudopneumoniae*. The cross-mapping remained minimal for all *Streptococcus* species outside the most closely related species. For example, while 43.6% of the mapped reads from the *S. pseudopneumoniae* genome mapped to the correct species, an additional 55.5% mapped to the close relatives *S. mitis, S. oralis*, and *S. pneumoniae*. Reads generated from genomes outside the genus *Streptococcus* generally did not map to the reference genome set (Fig. S2, Table S2). Across all reference genome set species and all samples simulated from non-*Streptococcus* spp. templates, the total mean coverage had an average of 0.02, which is within roundoff error and indicates that the presence of reads from other genera in metagenomic samples is unlikely to influence the results of mapping to this reference genome set.

### Metagenomic read mapping reveals taxon site-specificity and ecological relevance of reference genomes

To assess the distribution and abundance of streptococci across the oral cavity we used metagenomic short reads sequenced from oral samples and mapped them competitively to our selected oral *Streptococcus* spp. reference genomes. We mapped a total of 706 quality-filtered metagenomic samples containing 34.4 billion paired-end Illumina reads (Table S5). These samples had been collected from nine sites (buccal mucosa, keratinized gingiva, hard palate, tongue dorsum, throat, palatine tonsils, supragingival plaque, subgingival plaque, and saliva) in 144 volunteers and shotgun sequenced as part of the Human Microbiome Project (HMP) (Lloyd-Price et al. 2017). Using the data analysis platform anvi’o, we assessed the prevalence of genes and genomes within each sample, and we aggregated the data from genomes within the same species to generate species-level information.

Read mapping showed that each *Streptococcus* species preferentially colonized a subset of oral sites. In the heat map in Fig. 2A, each column represents an individual sample from the mouth and each row represents a species. Light colors indicate species making up the largest proportion of streptococci in a sample, whereas black indicates undetected species. Generally, the distribution of the streptococci in the buccal mucosa resembles their distribution in the keratinized gingiva (Fig. 2A). Their distribution in the tongue dorsum resembles their distribution in the throat and palatine tonsils, and their distribution in the supragingival plaque resembles their distribution in the subgingival plaque. This trend is common for oral taxa (Mager et al. 2003. Segata et al. 2012). The majority of the HMP samples come from three sites — buccal mucosa, tongue dorsum, and supragingival plaque — which represent the three major categories of host tissue found in the oral cavity: non-keratinized mucosa, keratinized mucosa, and enamel. Among these three sites, each species that was abundant enough for its distribution to be measured had several-fold greater relative abundance in one of these three sites than in the others.

**Figure 2:**
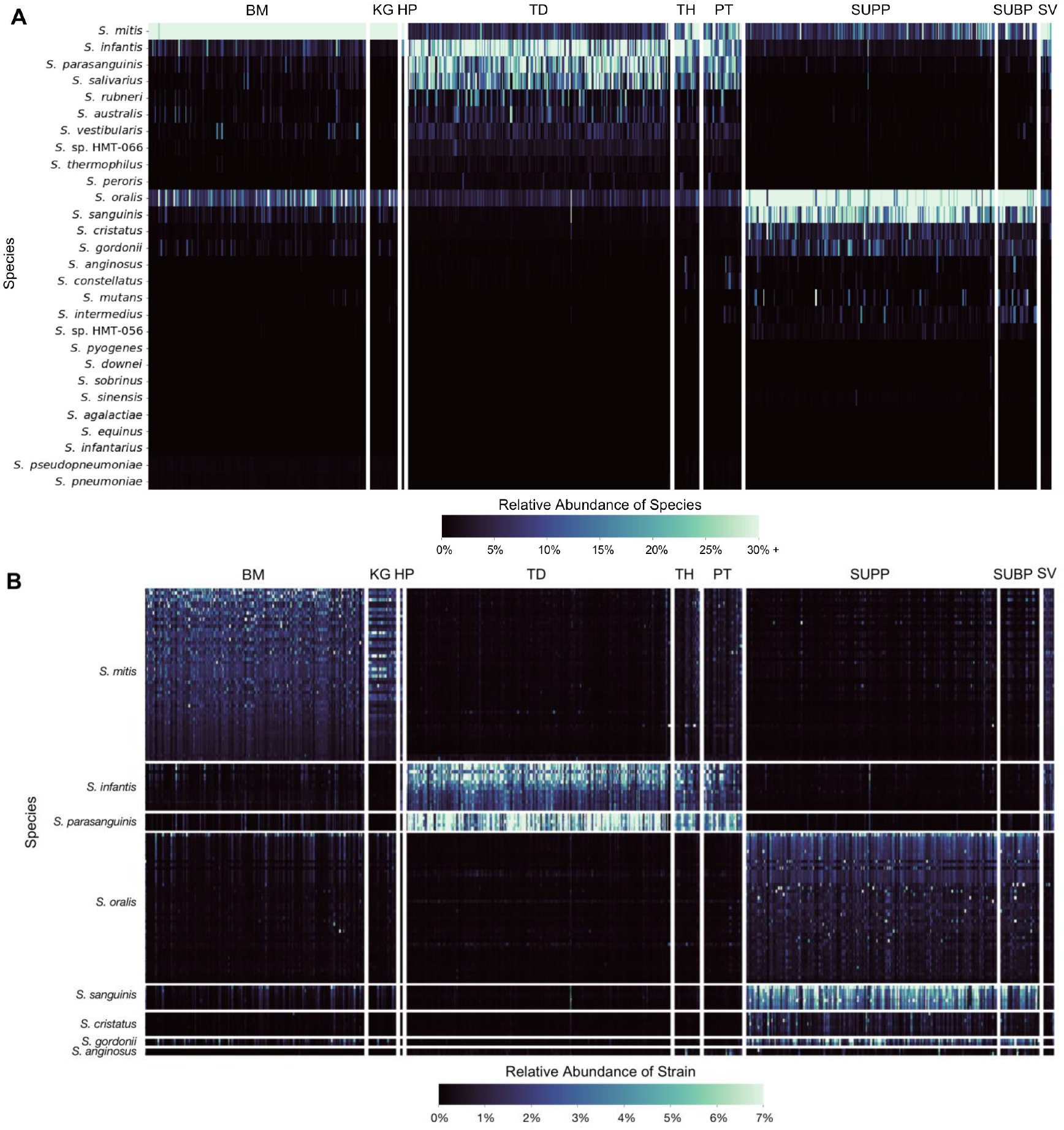
Mapping reveals differences in the abundance of species between oral sites and differences in the abundance of strains within an oral site. The heatmaps show the fraction of oral Streptococcus (A) species and (B) individual strains from species with more than one representative genome for each of the metagenomes sampled across nine oral sites. There are 183 buccal mucosa (BM), 23 keratinized gingiva (KG), 1 hard palate (HP), 220 tongue dorsum (TD), 21 throat (TH), 31 palatine tonsils (PT), 209 supragingival plaque (SUPP), 32 subgingival plaque (SUBP), and 8 saliva (SV) samples. The samples are grouped by site and then ranked by descending number of total reads. The strains are grouped first by species and then ranked by descending mean relative abundance across the site (BM, TD, or SUPP) where they are most abundant.

The closely related taxa *S. mitis, S. infantis*, and *S. oralis* showed distinct localization patterns. *S. mitis* was in high abundance on the buccal mucosa and keratinized gingiva, while *S. infantis* was abundant on the tongue, throat, and palatine tonsils and *S. oralis* was among the most abundant species in dental plaque. Thus, the higher resolution afforded by this whole-genome analysis made it possible to distinguish the mapping patterns of these taxa, which cannot be clearly resolved in analyses that rely on the 16S rRNA gene alone.

Whereas the results above provide species-level analysis by summing the reads that mapped to each of the representative genomes from a species, separating the mapping results for each reference genome shows that not all strains of a given species are equally represented in the oral cavity (Fig. 2B). Some *S. mitis, S. oralis*, and *S. infantis* reference genomes recruited more reads than others, indicative of greater similarity to the populations of these three species present in the sampled mouths (Fig. 2B). The differences in genome-level mapping indicate which sequenced genomes are most representative of the populations in the healthy mouth in these subjects. Overall, genomes within a species showed similar distribution patterns across the oral sites, providing no evidence for subspecialization to different sites within the named species.

Analysis of the breadth of coverage confirms differential site-specificity among closely related taxa by showing whether the gene content of a given sequenced genome matches the gene content of the population in the mouth. For each of the major oral streptococci, Fig. 3 indicates which of the ORFs in a representative genome were detected in 30 metagenomes from the three major oral sites. Among the closely-related *S. infantis, S. mitis*, and *S. oralis*, most of the ORFs in the genome were detected primarily in samples from a single habitat as indicated by the depth of coverage results: tongue dorsum for *S. infantis*, buccal mucosa for *S. mitis*, and supragingival plaque for *S. oralis* (Fig. 3). An additional pair of closely related taxa, *S. salivarius* and *S. vestibularis*, demonstrate more nuanced site-specificity: both taxa recruit many reads from tongue dorsum samples, but in *S. salivarius* more than 90% of ORFs were detected in most TD samples, whereas in *S. vestibularis* fewer than 75% were usually detected (Fig. 3, Table S7). This difference indicates that the gene content of *S. salivarius* is a good match to that of the tongue dorsum population: *S. salivarius* is a tongue specialist in the healthy oral microbiome. By contrast, *S. vestibularis*, which was originally isolated from the mucosa of the vestibule or the front of the mouth, reaches detection of >90% of ORFs in samples from buccal mucosa but not from tongue; it is apparently a sporadically abundant buccal mucosal taxon. The recruitment of reads by *S. vestibularis* from tongue samples is likely due to erroneous cross-mapping from the closely related and highly tongue-abundant *S. salivarius* (Table S2). Thus, data on breadth of coverage can reveal differential distributions among closely related taxa.

**Figure 3:**
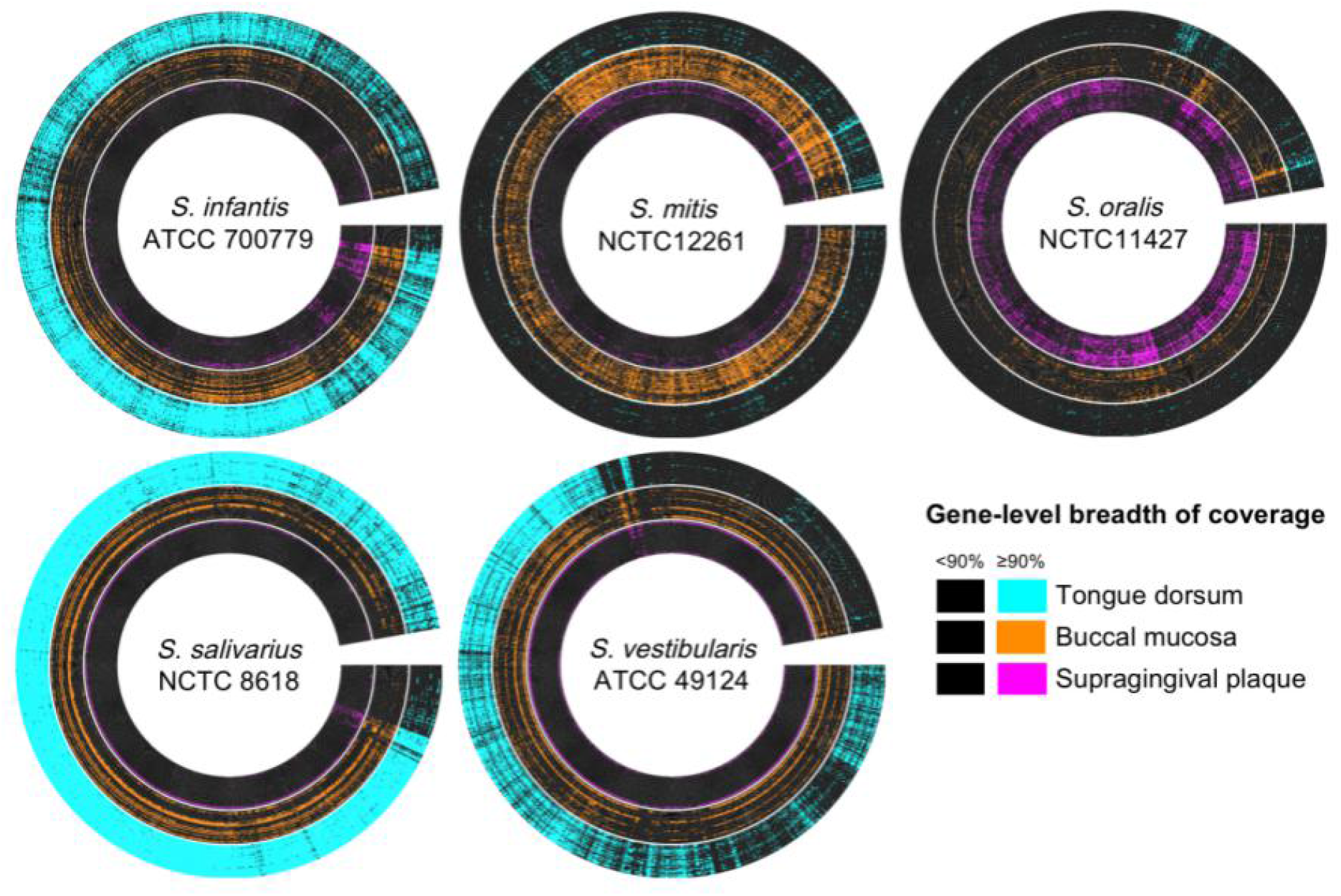
Breadth of coverage validates site tropisms and indicates how well the sequenced genome matches the gene content of the population in the mouth. The radial heatmap displays breadth the of coverage of the predicted open reading frames (ORF) from a representative genome from the 30 buccal mucosa, tongue dorsum, and supragingival plaque samples with the most quality-filtered reads. Each radius represents a predicted ORF. Each concentric ring represents a metagenomic sample. ORFs are black if their breadth of coverage is < 90% and color-coded by site if their breadth of coverage is ≥ 90%. Site tropisms of *S. infantis, S. mitis*, and *S. oralis* were confirmed, as most ORFs of S. infantis were detected in most samples from tongue dorsum, most ORFs of *S. mitis* were detected in samples from buccal mucosa, and most ORFs of *S. oralis* were detected in samples from supragingival plaque. Breadth of coverage indicates differing site tropisms of the closely related *S. salivarius* and *S. vestibularis*: most ORFs of *S. salivarius* were detected in tongue dorsum samples, while in *S. vestibularis*, most ORFs were detected in samples from buccal mucosa, and approximately a quarter of ORFs were not detected in samples from the tongue. The genomes displayed here represent the genomes from each species which recruited the most reads across all metagenomes and whose NCBI species matched our new species designations.

Whereas prior results using short regions of the 16S rRNA gene had suggested that a cluster of *Streptococcus* species contained oral generalists, genomic read mapping resolved this cluster into individual species that primarily localize to different sites. The diagram in Fig. 4A, modified from Mark Welch et al. 2019, shows habitat specialization based on oligotyping data in which the genus was divided into subsets, most of which were clusters of related species. The cluster containing *S. mitis, S. oralis*, and *S. infantis* appeared to be a generalist. Mapping shotgun sequencing reads identified a dozen *Streptococcus* species (Fig. 4B). These species include buccal mucosa specialist *S. mitis*, tongue dorsum specialists *S. infantis, S. australis, S.* sp. HMT-066, *S. parasanguinis, S. rubneri*, and *S. salivarius*, as well as supragingival plaque specialists *S. oralis, S. cristatus, S. gordonii*, and *S. sanguinis. S. vestibularis*, not shown in Fig. 4B, is potentially a buccal mucosa specialist despite its inclusion in the *S. salivarius* group in Fig. 4A. These data further show the importance of the species-level resolution provided by shotgun sequencing read mapping. Taken alone the 16S data classified the Mitis group as an apparent generalist taxon; however, the shotgun sequencing mapping data show that the species previously lumped into the Mitis group (*S. mitis, S. infantis, S. australis, S. oralis*, and *S. cristatus*) are specialists with preferences for either the buccal mucosa, tongue dorsum, or supragingival plaque.

**Figure 4:**
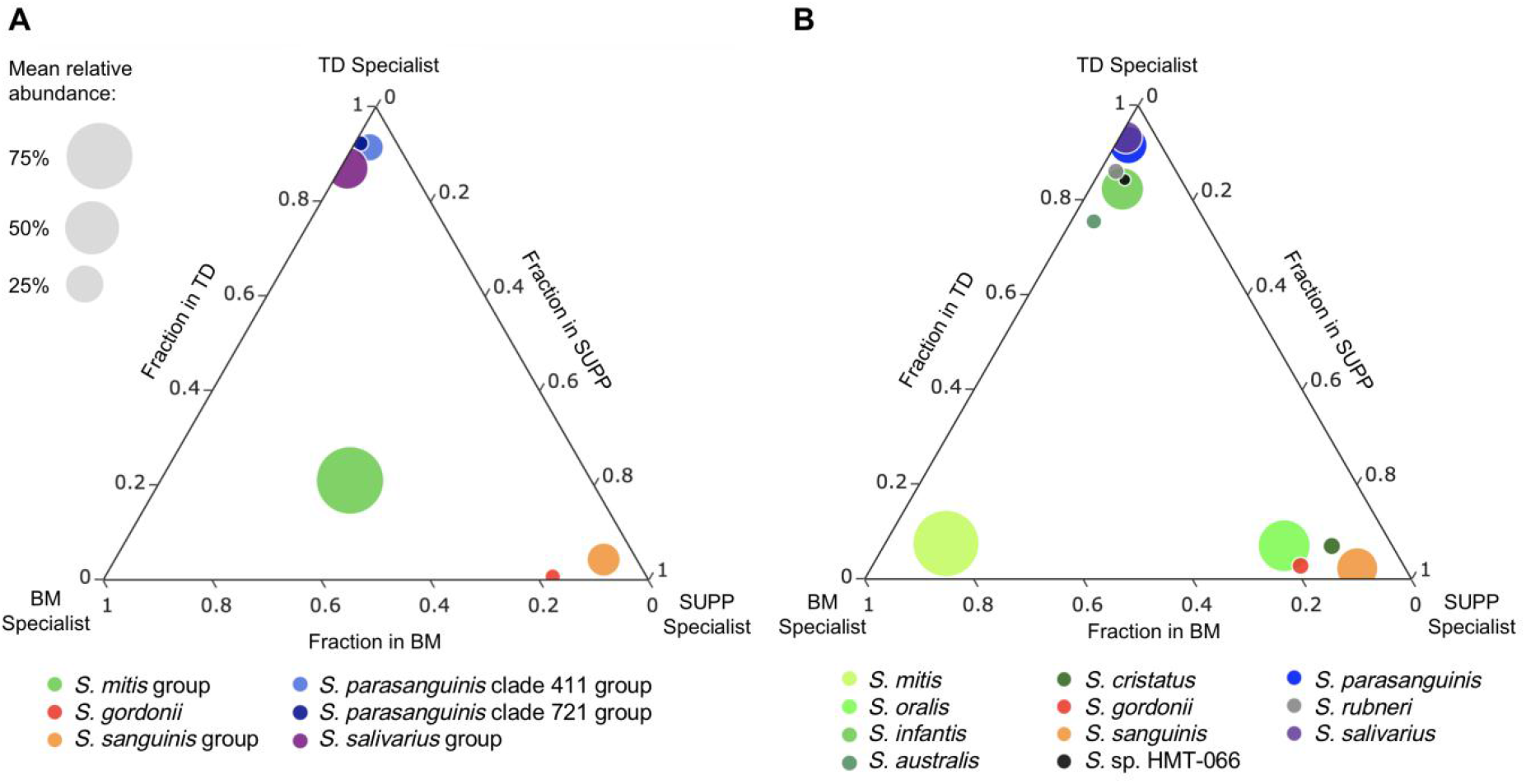
Mapping of whole-genome sequence data reveals site-specificity of species that cannot be distinguished based on 16S sequencing data. These ternary plots show the mean relative abundance of streptococci across three oral sites – buccal mucosa (BM), tongue dorsum (TD), and supragingival plaque (SUPP) – estimated via (A) oligotyping analysis of 16S gene sequencing data using the V1-V3 region of the 16S rRNA gene and (B) mapping whole-genome shotgun sequencing reads to the reference genome set. Bubbles are color-coded by taxon. Species in B and the groups they were lumped into in A are shown in different hues of the same color. Bubble size is proportionate to the mean relative abundance in the oral sites where the taxon is most abundant. The fraction of mapping to a site is calculated by dividing the mean relative abundance for a taxon in that site by the sum of the mean relative abundances of that taxon in all three sites. The bubbles of site-specialist taxa cluster near a corner of the plot. A is adapted from Mark Welch et al. 2019 and based on data from Eren et al. 2014. Species with a mean abundance ≥ 2% averaged across all samples from at least one site (excluding S. vestibularis) are shown in B.

### Functional annotation of *S. mitis, S. oralis*, and *S. infantis* reveals no species-specific core functions that could drive localization to different sites

Pangenomics, which entails the identification of essential core and nonessential accessory genes for a set of related microbial genomes, can be used to identify genes involved in adaptation to distinct microhabitats that may give rise to the spatial distribution patterns revealed by metagenomics (Scholz et al. 2016, Nayfach et al. 2016, Delmont and Eren 2018). Seeking to identify genes underlying these differential distribution patterns, we constructed a pangenome of the genus *Streptococcus* using the reference genome set generated above. The visualization of the pangenome shows the genomes hierarchically clustered to group together the genomes with more similar gene content (Fig. 5). This clustering, based on the prevalence in each genome of the 18,895 gene clusters of the pangenome, gave results that were broadly consistent with the phylogenomic tree: both analysis methods grouped the same genomes into species-level clusters with multiple genomes and placed the *S. pneumoniae* and *S. pseudopneumoniae* genomes within the *S. mitis* clade and the *S. peroris* genome within the *S. infantis* clade (Fig. S3). A set of 606 core gene clusters, constituting 27-38% of the clusters in each genome, was found across all the reference genomes (Fig. 5).

**Figure 5:**
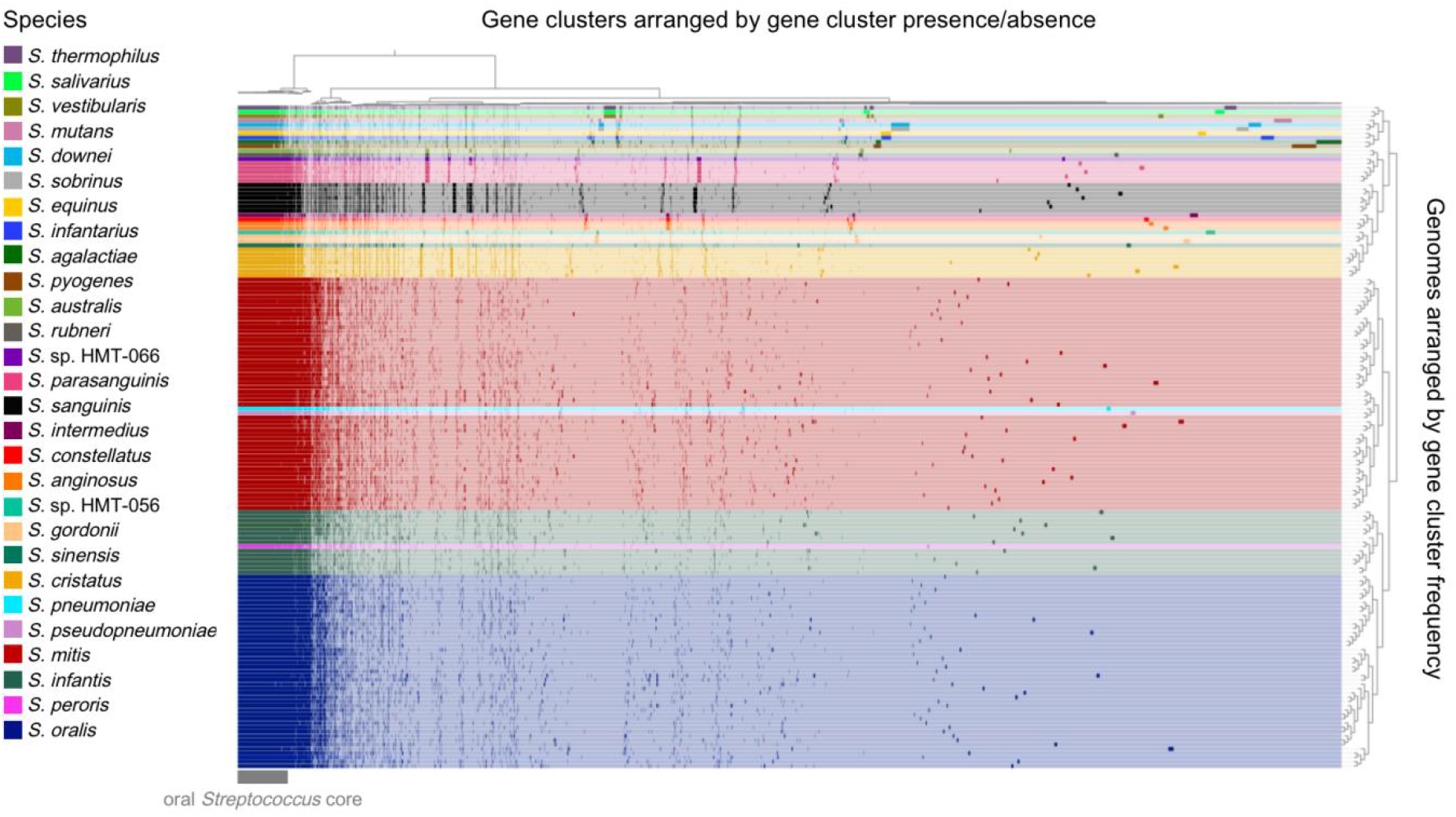
The human oral cavity has numerous endemic streptococci species, both homogeneous and heterogeneous, that possess many shared core genes. The genomes are clustered by the frequencies of their gene clusters and color-coded by species. The gene clusters are clustered according to their presence or absence in each genome; presence is denoted by a dark shade and absence by a light shade of the color representing each species. The set of 154 oral streptococci genomes includes representative genomes with shared ANI values of < 95%, so species with more genomic nucleotide-level diversity have a larger number of representative genomes. The number of representatives is also affected by the availability of genomes for a species. The 18,895 distinct gene clusters of the pangenome include 606 core genes that occur in every genome. Large sets of species-specific core genes distinguish several of the Streptococcus species.

Although some *Streptococcus* species in the pangenome possess large blocks of species-specific gene clusters, others – notably *S. mitis, S. oralis*, and *S. infantis* – do not. Inspection of the pangenome shows blocks of gene clusters characteristic of individual species such as *S. cristatus, S. sanguinis*, and *S. parasanguinis*, as well as blocks characteristic of groups of closely related species such as *S. salivarius, S. vestibularis*, and *S. thermophilus* (Fig. 5). *S. sanguinis*, for example, has a well-defined block of species-specific core genes that account for 2.3-2.5% of the gene clusters in its genome. By contrast, and consistent with the results of a similar pangenome constructed by Velsko et al. (2019), *S. mitis*, *S. oralis*, and *S. infantis* appear to share many accessory genes in common but do not have major blocks of gene clusters unique to each species. The apparent similarity of the species-specific core for these three species contrasts with the observed differences in their distribution.

Constructing a targeted pangenome with only *S. mitis, S. infantis*, and *S. oralis* indicated that there are almost no annotated functions that were unique and core to one of the species to the exclusion of the others. In the targeted pangenome, constructed using a more stringent value of the “minbit” parameter for eliminating clusters with low amino acid sequence similarity, modest blocks of species-specific core genes were present for each of *S. mitis, S. oralis*, and *S. infantis* (Fig. 6). *S. mitis* and *S. oralis* also shared a block of core genes, while *S. infantis*, shared two core gene clusters with *S. oralis* and one with *S. mitis*. To determine whether these gene clusters had unique and potentially niche-defining functions, we carried out functional annotation of each called gene using the Pfam, NCBI COG, and eggNOG databases. The results indicated that nearly all the species-specific core gene clusters were annotated with functions found in all three species. Depending on the annotation source, each species had between zero and two annotated functions that were both unique and core to the species. No annotated function was unique and core to two species. Thus, use of a more stringent clustering parameter reveals a set of species-specific core genes for each taxon but these are distinguished by amino acid divergence and not by functional divergence, as nearly as can be discerned using current annotation databases.

**Figure 6:**
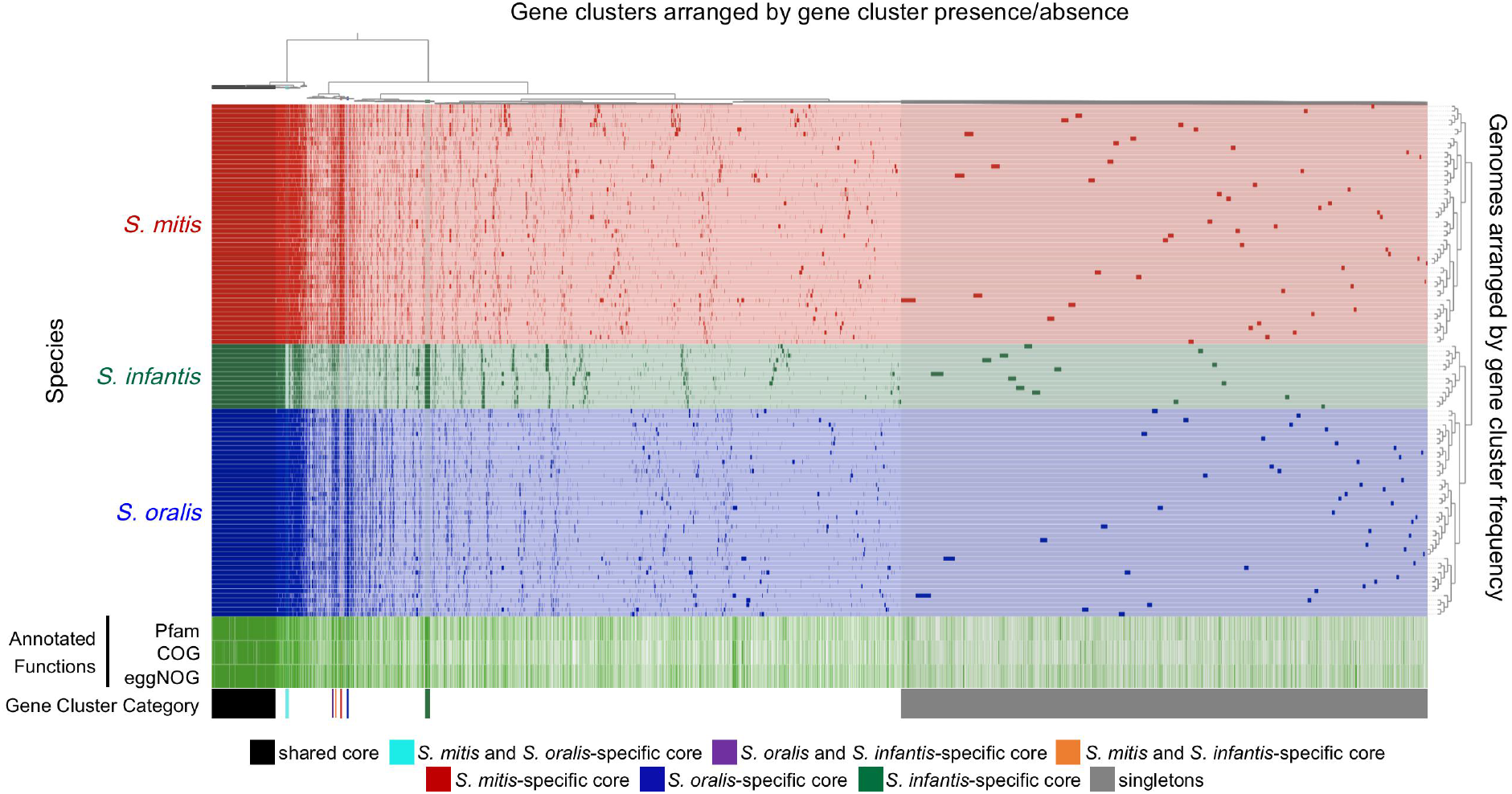
Nearly all genes that are core to the individual species *S. mitis, S. oralis*, and *S. infantis* are found in all three species and detected across habitats. Genes and genomes are clustered as in Fig. 5. Gene clusters present in all genomes are marked shared core. Gene clusters present in all genomes of one or two species and none of the genomes of the other species are marked as species-specific core. Gene clusters present in a single genome are marked as singletons. The “Annotated Functions” layers indicate whether the gene cluster has (green) or has not (white) been assigned a Pfam, COG, or eggNOG function.

## Discussion

For complex microbial populations composed of numerous closely-related species and strains, genome-scale analysis provides the resolution necessary to demonstrate site-specificity. The idea that most species of oral bacteria are site-specialists originated in culture-based studies extending back to the 1960s (Gibbons et al. 1963, Gibbons et al. 1964 A, Gibbons et al. 1964 B, Gordon and Gibbons 1966, Gordon and Jong 1968, Frandsen et al. 1991) and was strengthened by the systems-level view provided by culture-independent studies in which total DNA is extracted from the samples and analyzed for the presence of microbes via checkerboard DNA-DNA hybridization (Mager et al. 2003) or sequencing of the 16S ribosomal RNA gene (Aas et al. 2005, Zaura et al. 2009, Human Microbiome Project Consortium 2012, Huse et al. 2012, Peterson et al. 2013, Mark Welch 2014, Eren et al. 2014, Hall et al. 2017, Bernardi et al. 2020). Using the greater resolution provided by mapping whole-genome sequencing data, we could determine the localization for *Streptococcus* species and assess how the site-specialist hypothesis applied to closely related species, not distinguishable by their 16S rRNA gene sequence. We determined that all the oral *Streptococcus* species that could be detected by short-read mapping were site-specialists. *S. sanguinis, S. cristatus, S. gordonii*, and *S. mutans* were among the most abundant in the supragingival plaque, replicating prior findings based on 16S sequencing data (Huse et al. 2012, Peterson et al. 2013, Eren et al. 2014, Hall et al. 2017). *S. australis, S. salivarius*, and *S. parasanguinis* were among the most abundant species on the tongue dorsum, likewise replicating the findings of 16S studies (Aas et al. 2005, Mark Welch et al. 2014, Eren et al. 2014, Bernardi et al. 2020). Whole-genome sequencing data differentiated between *S. mitis, S. oralis*, and *S. infantis*, which could not be distinguished in 16S rRNA gene studies (Zaura et al. 2009, Huse et al. 2012, Mark Welch 2014, Eren et al. 2014, Hall et al. 2017). When the attempt was made to distinguish between these species, *S. oralis* and *S. infantis* were either scarce or undetected (Aas et al. 2005, Peterson et al. 2013). Our results indicated that these three closely related species preferentially localized to different sites. 16S rRNA gene studies could not differentiate between *S. salivarius* and *S. vestibularis* and indicated that OTUs containing both species have the greatest relative abundance in the tongue dorsum (Zaura et al. 2009, Huse et al. 2012, Mark Welch et al. 2014, Eren et al. 2014, Hall et al. 2017). While cross-mapping makes distinguishing between the two species with shotgun reads difficult as well, the gene-level mapping data indicate *S. salivarius* is a tongue dorsum specialist whereas *S. vestibularis* is a buccal mucosa specialist. The distribution of both groups of closely related species demonstrates that taxonomy is not always a clear indicator of the spatial niche where oral species specialize. Our analysis indicates *S. rubneri* and *S*. sp. HMT-066, more recently identified species not included in earlier studies, are tongue dorsum specialists.

The challenge of distinguishing between *S. salivarius* and *S. vestibularis* illustrates the limitations of mapping shotgun sequencing data. Issues with the interpretation of short reads mapped to reference genomes arise due to stretches of high homology that occur in closely related taxa or can be generated by mobile elements, horizontal transfer, or genes highly conserved at the nucleotide level such as ribosomal RNA genes. Thus, short reads have a certain probability of mapping to related genomes, or to unrelated genomes that share a particular mobile element. To account for these issues when interpreting mapping data, one must assess whether the mapped reads are concentrated in small regions of the genome or whether they cover the expected fraction of the genome given their abundance (Martin et al. 2012, Kraal et al. 2014). One approach to mitigate the cross-mapping problem is profiling the abundance of taxa using only read recruitment to taxon-specific core genes (Liu et al. 2011, Segata et al. 2012, Sunagawa et al. 2013, Truong et al. 2015). However, this approach is best suited for analyzing taxa that have large complements of unique core genes, unlike the oral *Streptococcus* spp. As with other techniques, rare taxa present a challenge for short-read mapping. Reliable detection of a taxon requires its reads to be sufficiently represented in a metagenome to cover the taxon core, which is not always the case for low abundance taxa. Despite these challenges, short-read mapping results interpreted with appropriate care provide unparalleled insights into the composition and functional potential of native microbial communities in the human mouth.

Although *S. mitis*, *S. oralis*, and *S. infantis* were each abundant at one primary site, each was also detected in a subset of samples at the other sites. Detection of site-specialists outside their favored sites could indicate the presence of a strain or sub-population with a site specialization different from the rest of the species. While the species *Haemophilus parainfluenzae* contains strains that apparently specialize to different sites (Utter et al. 2020), the mapping results for individual strains indicate this is not true of oral streptococci. Instead, detection of species like *S. mitis* or *S. infantis* outside their primary sites might be due to the colonization of favorable microhabitats within unfavorable oral sites. For example, the supragingival plaque biofilm is heterogeneous and contains various complex structures (Mark Welch et al. 2016) in which specialized microhabitats for otherwise rare oral microbes may exist. A variety of habitats may also be created by temporal succession, as the abrasion of the tooth surface and the shedding of old host cells would be expected to create fresh substrate for new biofilm formation, creating a shifting mosaic steady state in which supragingival plaque in both the initial and the late stages of successional development coexist. *S. mitis* is a primary colonizer of tooth surfaces and is abundant in new plaque (Nyvad and Kilian 1990, Frandsen et al. 1991, Li et al. 2004); however, as dental plaque matures it begins to be supplanted by other taxa (Ramberg et al. 2003). Our detection of low abundances of *S. mitis* in the supragingival plaque samples may correspond to detection of *S. mitis* in patches of initial plaque. The low abundance of cells that primarily localize to other sites may also correspond to detection of bacterial sojourners, bacteria deposited at the site where the conditions are unfavorable for colonization and growth. Finally, our conclusions about the distribution of *Streptococcus* species are based on our analysis of metagenomic samples, which may have been biased by sampling methodology (McInnes and Cutting 2010). Cells are shed into the saliva from all oral sites and dispersed throughout the oral cavity by salivary flow. Because the HMP sampling protocols do not include precautions to exclude saliva, the samples may include cells shed from other sites.

The distribution of the detectable *Streptococcus* species suggests specialized adaptation for different spatial niches within the oral cavity. The development of pangenomic and metapangenomic approaches to studying community ecology has increased the ease of identifying repertoires of unique genes associated with the phenotypes underlying niche adaptation. Pangenomics has been used to identify genes differentiating the *Streptococcus* communities associated with hosts from different species (He et al. 2017, Kawaski et al. 2018, Richards et al. 2019) and geographic locations (Kayansamruaj et al. 2015, Iversen et al. 2020). Similar techniques have identified species-specific core genes that distinguish various *Streptococcus* species (Lefébure and Stanhope 2007, Croxen et al. 2018, Zheng et al. 2017, Velsko et al. 2019, Iversen et al. 2020, Gonzales-Siles et al. 2020). To better understand the adaptation of *Streptococcus* species to different spatial niches within the mouth, we searched for genes that differentiated the closely related species *S. mitis, S. oralis*, and *S. infantis* using pan- and metapangenomics. Yet, we found nearly no species-specific core genes with unique functions that might explain their spatial distribution. One limit of this analysis is the limits to the accuracy and specificity of the tools presently available for functional annotation. Some of the core gene clusters specific to one or two species were not annotated, while others received annotations that were either vague or based on functions characterized for proteins from taxa as distant as eukaryotes.

One phenotype that would be reasonably expected to distinguish the three species is the capacity to adhere to different substrates. To resist the shearing force of salivary flow and remain in a preferred environment, non-motile streptococci must adhere to that site. Oral streptococci possess many adhesins that mediate highly specific adhesion to components of the acquired salivary pellicle, extracellular matrix, host cells, and other microbes (Nobbs et al. 2009), yet none of the species-specific core genes were annotated with functions suggesting a role in adhesion. Thus, phenotypic differences between the species may also be due to factors beyond the prevalence of protein-coding genes — like small sequence differences in conserved genes, differences in gene expression, or differences in gene copy number. Lefébure and Stanhope (2007) previously found that numerous core genes shared between several other *Streptococcus* species, especially those related to colonization and biofilm formation, were subject to positive selection, supporting the idea that differences between protein-coding genes sharing similar functions may contribute to niche adaptation.

While we were unable to identify potential drivers of the different spatial distribution patterns displayed by *S. mitis, S. oralis*, and *S. infantis* with an ‘omics-based approach, the underlying mechanisms could also be investigated using methods drawn from biochemistry or molecular biology. Binding assays could be used to identify proteins responsible for interspecies differences in adhesion to substrates or congregation with other taxa present at the oral sites. Knock-out mutant generation and change of function assays could be used to identify the genes more broadly associated with differences in site-specificity. One barrier to such approaches is the lack of in vitro models that replicate the complexity of the oral environments as each oral site contains numerous substrates, chemical gradients, and taxa arranged into structures of varying complexity.

Strains of *S. mitis, S. oralis*, and *S. infantis* possess unusually high divergence from one another, as measured by ANI, but our results support the idea that the species as currently defined are biologically meaningful. While there is not a standard prokaryotic species definition, bacterial species are generally considered to consist of collections of strains that are genomically coherent; they share a greater gene content and sequence similarity with each other than with other species (Konstantinidis and Tiedje 2005). Intraspecies genomic coherence is maintained through gene exchange, and barriers to recombination have been proposed as the limits to bacterial and archaeal species (Bobay and Ochman 2017). A genomic distance of around 95% ANI is often recommended as a species boundary as this similarity score circumscribes most recognized species (Konstantinidis and Tiedje 2005, Jain et al. 2018, Olm et al. 2020, Park et al. 2020). However, the members of multiple recognized *Streptococcus* Mitis group species share mean ANIs between 90% and 95% (Jensen et al. 2016). When Park et al. (2020) proposed a new taxonomy for the RefSeq genomes in the Genome Taxonomy Database (GTDB) using a 95% ANI species boundary, they subdivided these species into as many as 50 species clusters. Our phylogenomic analysis supports the idea that the current named oral *Streptococcus* species are genomically coherent, including the genomically divergent species — *S. mitis, S. oralis*, and *S. infantis.* The *Streptococcus* spp. genomes we analyzed formed distinct clusters with respect to ANI that corresponded to existing species classifications. In addition to genomic coherence, biologically meaningful species are expected to share consistent phenotypes (Konstantinidis and Tiedje 2005). Mapping indicated that members of a named species shared a common localization phenotype, which differed between closely related species like *S. mitis*, *S. oralis*, and *S. infantis* or *S. salivarius* and *S. vestibularis*. These results support the validity of the recognized oral streptococci species and highlight the difficulty of selecting a universal genomic similarity threshold to circumscribe all prokaryote species.

## Materials and Methods

We used a workflow adapted from Delmont and Eren (2018) to perform metapangenomic analyses in the anvi’o v7 platform (Eren et al. 2021) with Python v3.7.9. All shell scripts used in our analyses are provided in the supplemental material.

### Reference Genomes and Metagenomes

Unbiased metagenomic read mapping requires a set of reference genomes to which the reads will be aligned that are accurately identified and representative of the range of diversity present in the taxa of interest. The inclusion of a larger number of representative genomes for some taxa than for others would bias read recruitment in favor of the better-represented taxa. Therefore, following previous authors (Delmont et al. 2018, Almeida et al. 2019, Olm et al. 2021) we restricted the set of reference genomes to those that shared no more than a given percentage of average nucleotide identity (ANI), in this case 95%. We used RefSeq genomes from the named *Streptococcus* species and unnamed *Streptococcus* “human microbial taxa” (HMT) in the eHOMD (http://www.homd.org). We also included genomes sequenced from human isolates if there was evidence of their presence in the human oral cavity (Tetz et al. 2019, Shen et al. 2002, Huch et al. 2013, Bernardi et al. 2020). The GTDB groups RefSeq genomes into clusters sharing ≥ 95% ANI (Parks et al. 2020). We chose one representative from each group (Table S1) that had a completeness of ≥ 90% estimated by CheckM (Parks et al. 2014). When choosing representatives, we also preferentially selected type strains and strains available from culture collections as well as genomes with high completeness and low contamination scores estimated by CheckM. Where possible within these constraints, we chose the representative genome identified by the GTDB. We added two additional genomes for eHOMD human microbial taxa (HMT) that were sequenced after the creation of the GTDB and substituted two GTDB cluster representatives for more recently sequenced genomes from the same strain that was more complete.

We downloaded the metagenomes used in this study from the Human Microbiome Project (HMP) Data Portal. These metagenomes consisted of 101-bp paired-end reads sequenced from samples collected from nine oral sites in phases I and II of the HMP. We downloaded all metagenomes uploaded through 12/20/2016 for oral sites that had at least 100 samples uploaded through this date and downloaded all metagenomes uploaded through 6/1/2021 for the other sites.

### Data Cleaning

We used the anvi’o program ‘anvi-compute-genome-similarity’ to calculate the ANI between all the genomes and cluster them based on these ANI values. This script used the program pyANI and the ANI BLAST algorithm (Pritchard et al. 2016). The *S. mitis* 4928STDY7071560 genome (GCF_902159415.1) was eliminated from the reference genome set as it shared no more than 85% ANI with any other *Streptococcus* spp. genome. *S. periodonticum* KCOM 2412 (GCF_003963555.1) was eliminated because it shared an ANI of > 95% with the *S. anginosus* type strain sequence, *S. anginosus* NCTC10713 (GCF_900636475.1). To avoid downstream problems, contigs smaller than 200 nucleotides were dropped from the reference genomes with the ‘anvi-script-reformat-fasta’ and all IUPAC ambiguity codes were replaced with “N”s.

Before the genomes were made publicly available, likely human reads had been removed from the samples. We performed additional quality-filtering of the metagenomic reads using ‘iu-filter-quality-minoche’ (available from https://github.com/merenlab/illumina-utils) a program that implements the recommendations of Minoche, Dohm, and Himmelbauer (2011) for improving the quality of Illumina sequencing data (Eren et al. 2013).

### Reference Genome Annotation

With ‘anvi-gen-contigs-database,’ we identified ORFs using a k-mer size of 4 and Prodigal v2.6.3 (Hyatt et al. 2010). First, we used ‘anvi-run-hmms’ to search for Hidden Markov Models (HMMs) against anvi’o’s four default HMM profiles using hmmscan from HMMER v3.2.1 (Eddy, 2009). Then, we used ‘anvi-run-pfams’ to match gene clusters with functions from the European Bioinformatics Institute’s Pfam database with hmmsearch from HMMER v3.2.1. Finally, we used ‘anvi-run-cogs’ to match gene clusters with functions from the updated 2020 version of NCBI’s Clusters of Orthologous Groups database (Tatusov et al. 2000) with NCBI’s Protein-Protein BLAST v2.10.1+ (Altschul et al. 1990). We annotated amino acid sequences, which were exported from the contigs database with ‘anvi-get-sequences-for-gene-calls,’ using eggNOG-mapper v2 with precomputed eggNOG v5 clusters through the online interface (http://eggnog5.embl.de/#/app/emapper) and imported the annotations into the contigs database with ‘anvi-script-run-eggnog-mapper’ (Huerta-Cepas et al. 2017, Huerta-Cepas et al. 2019). For each source, the function most frequently annotated for the amino acid sequences in that gene cluster was considered the representative function for the gene cluster.

### Phylogenomics

To check the genomes’ NCBI species designations, we used ‘anvi-gen-phylogenomic-tree’ and FastTree v2.1.3 SSE3 (Price et al. 2010) to generate a phylogenomic tree with the *Streptococcus* spp. reference genomes and a *Lactobacillus crispatus* genome included as an outgroup. The tree was based on the amino acid sequences of 205 single-copy core genes present in all 154 *Streptococcus* spp. genomes acquired with ‘anvi-get-sequences-for-gene-clusters’ and aligned with MUSCLE. To differentiate between *S. mitis, S. pneumoniae*, and *S. pseudopneumoniae* genomes, we aligned *S. pneumoniae* and *S. pseudopneumoniae* species-specific marker sequences identified by Croxen et al. (2018) to all the genomes with BLASTn (Zhang et al. 2000).

### Metagenomics

To assess the representation of the oral streptococci in the metagenomes, we mapped the metagenomes to the reference genome set. To reduce non-specific mapping, we competitively mapped the reads to the reference genomes set with bowtie2 v2.4.1 (Langmead and Salzberg, 2012), so each read was only mapped to the one genome that provided the closest match. Using bowtie2, we first generated a reference index for mapping and then mapped the reads to the genome set using bowtie2 v2.4.1 with the “--very-sensitive,” “--end-to-end,” and “--no-unal” flags. We used Samtools v1.9 (Li et al. 2009) to sort and index the read alignment data generated by bowtie2. Using ‘anvi-single-profile,’ we used the BAM files output by Samtools to create an anvi’o single-profile database for each metagenome’s alignment data. With ‘anvi-merge-profile,’ we merged the single-profile databases for all metagenomes. We calculated mapping metrics for the reads aligned to each genome from each metagenome with ‘anvi-summarize.’ To quantify the mapping results, we first averaged the depth of coverage (the number of reads mapped to a nucleotide position) across each of the reference genomes. Then to measure the relative abundance of one species among the oral Streptococci, we divided the total mean depth of coverage for a species’ genomes by the total mean depth of coverage for all the reference genomes.

### Pangenomics

To evaluate the distribution of genes within and between the human oral *Streptococcus* species, we used ‘anvi-pan-genome’ to construct an anvi’o pangenome database from the annotated reference genomes. This program first used Protein-Protein BLAST v2.10.1+ to find similar gene calls throughout all the genomes and used MUSCLE v3.8.425 (Edgar, 2004) to align the genes. The gene calls were clustered based on the homology of their translated amino acid sequences with the MCL algorithm using an MCL-inflation parameter of 10 while weak matches were eliminated using a minbit heuristic of 0.5 (Van Dongen and Abreu-Goodger, 2012). Finally, the genomes were hierarchically clustered based on the frequencies of the gene clusters they contained using Euclidean distances with Ward’s method, and the gene clusters themselves were hierarchically clustered based on their presence or absence within the genomes using Euclidean distances with Ward’s method. We used ‘anvi-compute-functional-enrichment’ to calculate the fraction of the genomes from each species annotated with that function and to select a representative function from each of the three annotation sources for each gene cluster based on which function was annotated most frequently. We created a more targeted pangenome with just the *S. mitis, S. oralis*, and *S. infantis* genomes as above, except we used a minbit heuristic of 0.8 due to the narrower taxonomic scope of this pangenome.

## Acknowledgments

We would like to thank Floyd Dewhirst and A. Murat Eren for helpful discussions and Rich Fox for expert systems administration and assistance with using the Josephine Bay Paul Center servers at the Marine Biological Laboratory. This work was supported by NIH NIDCR R01 DE 022586 (G.G.B.) and NIH NIDCR R01 DE 027958 (J.M.W.).

## Author Contributions

A.R.M., J.T.M., G.G.B., and J.M.W. designed the study. A.R.M. performed the analysis. A.R.M. and J.M.W. drafted the manuscript. All authors critically revised the manuscript and approved the final version.

## Supplemental Materials

**Figure S1:**
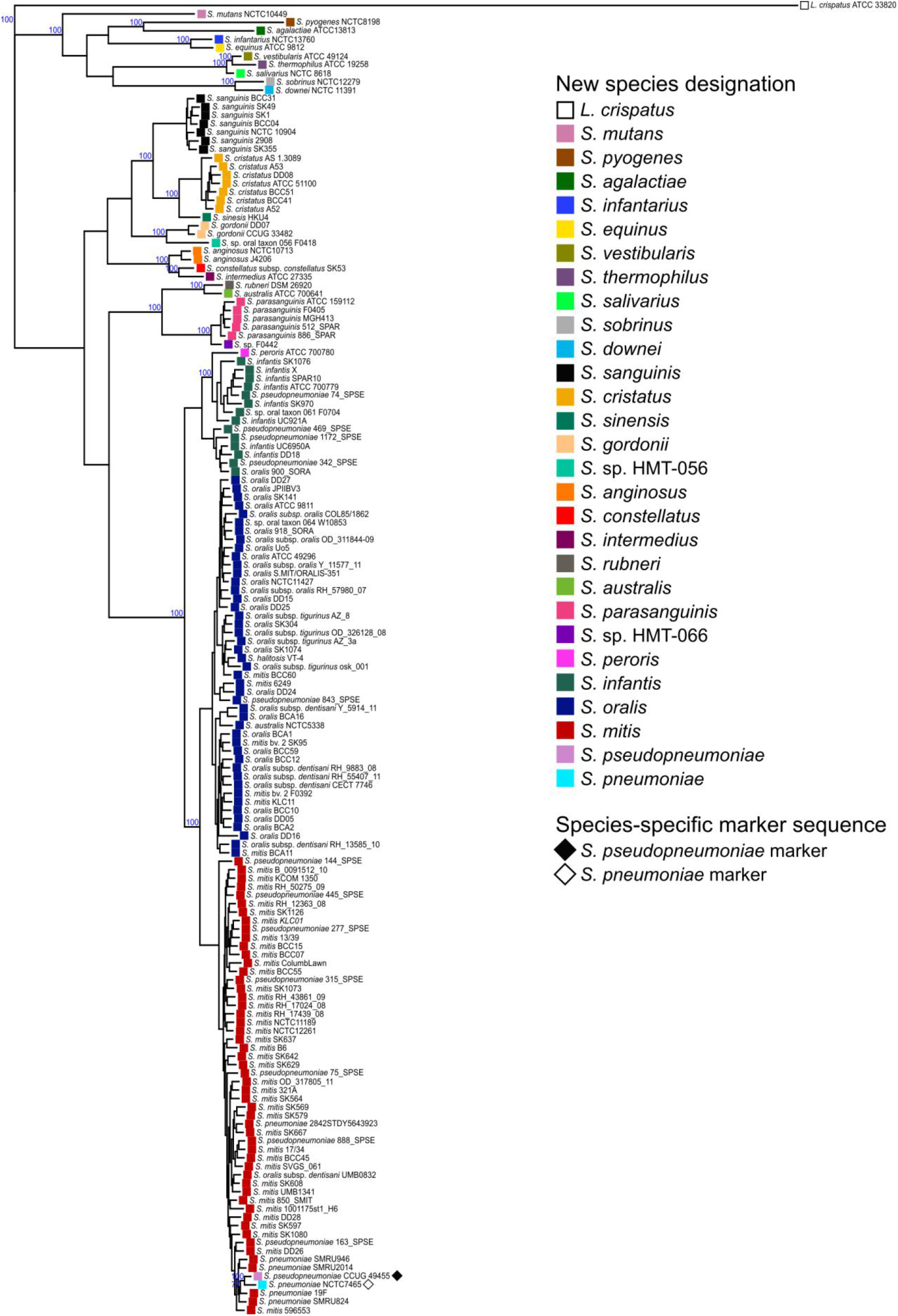
A phylogenomic tree based on 205 single-copy core genes (SCGs) indicates many of the mitis group reference genomes have incorrect NCBI species designations. The small text to the right of each node indicates the NCBI taxonomic designation of each genome. The colored labels indicate the revised species designation assigned to the genome. A “♦” indicates that the genome contains a > 99% identity match for the *S. pseudopneumoniae* marker genes and a “♢” indicates that the genome contains a > 99% identity match for the *S. pneumoniae* marker genes. Nodes that delineate species clusters are annotated with blue bootstrap values.

**Figure S2:**
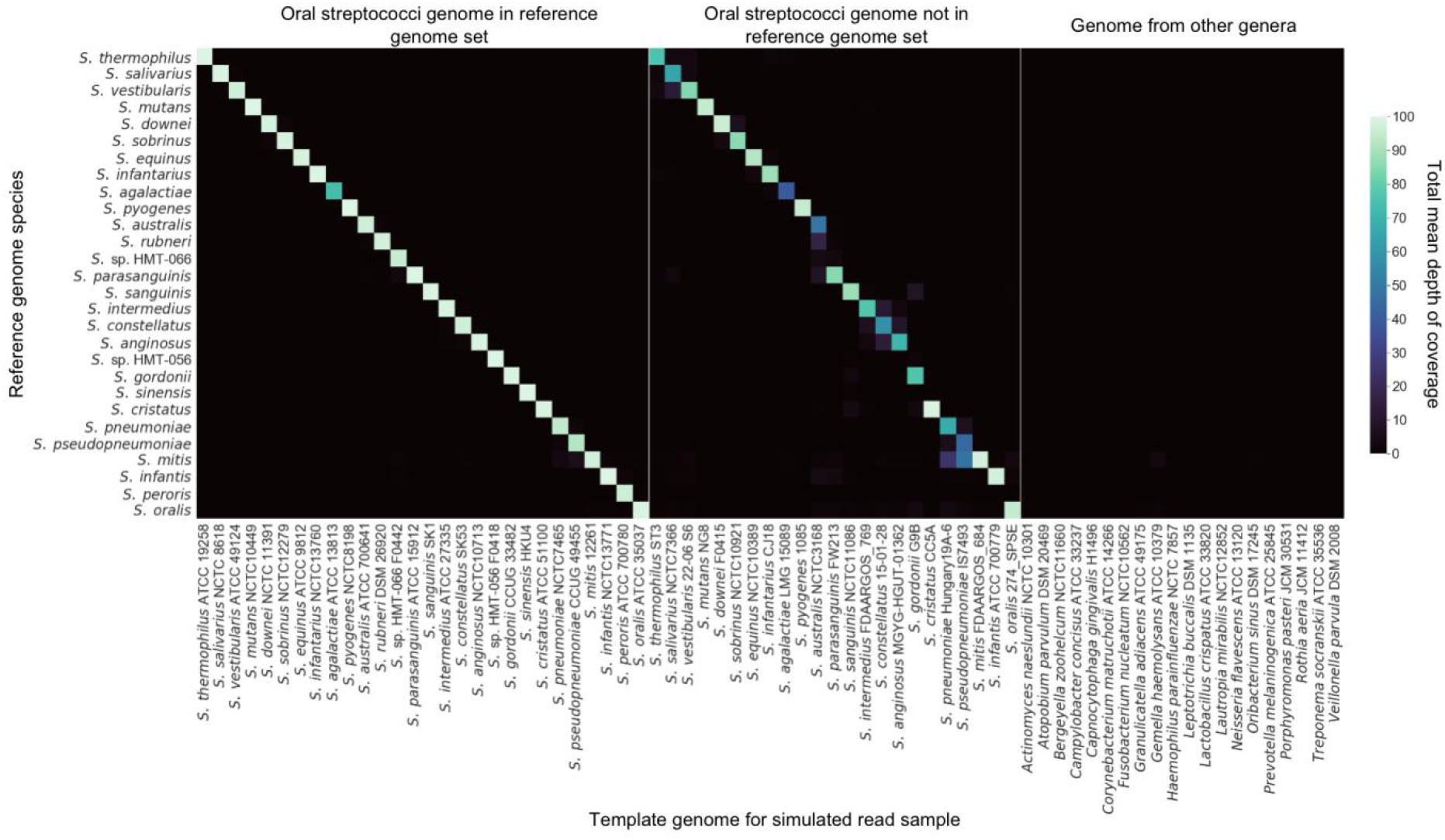
Simulated reads map with a high degree of specificity. For each simulated read sample, the matrix displays the sum of the mean depth of coverage values for all reference genome with the same species. The depth of coverage was average across all nucleotide positions in a reference genome. The reference genome species are arranged by their approximate order in the pangenome. The simulated samples are grouped into reads simulated from streptococci sequences in the reference genome set, streptococci sequences not in the reference genome set, and sequences from other major oral genera. Within the first two groups, the samples are arranged by the order of their species in the pangenome.

**Figure S3:**
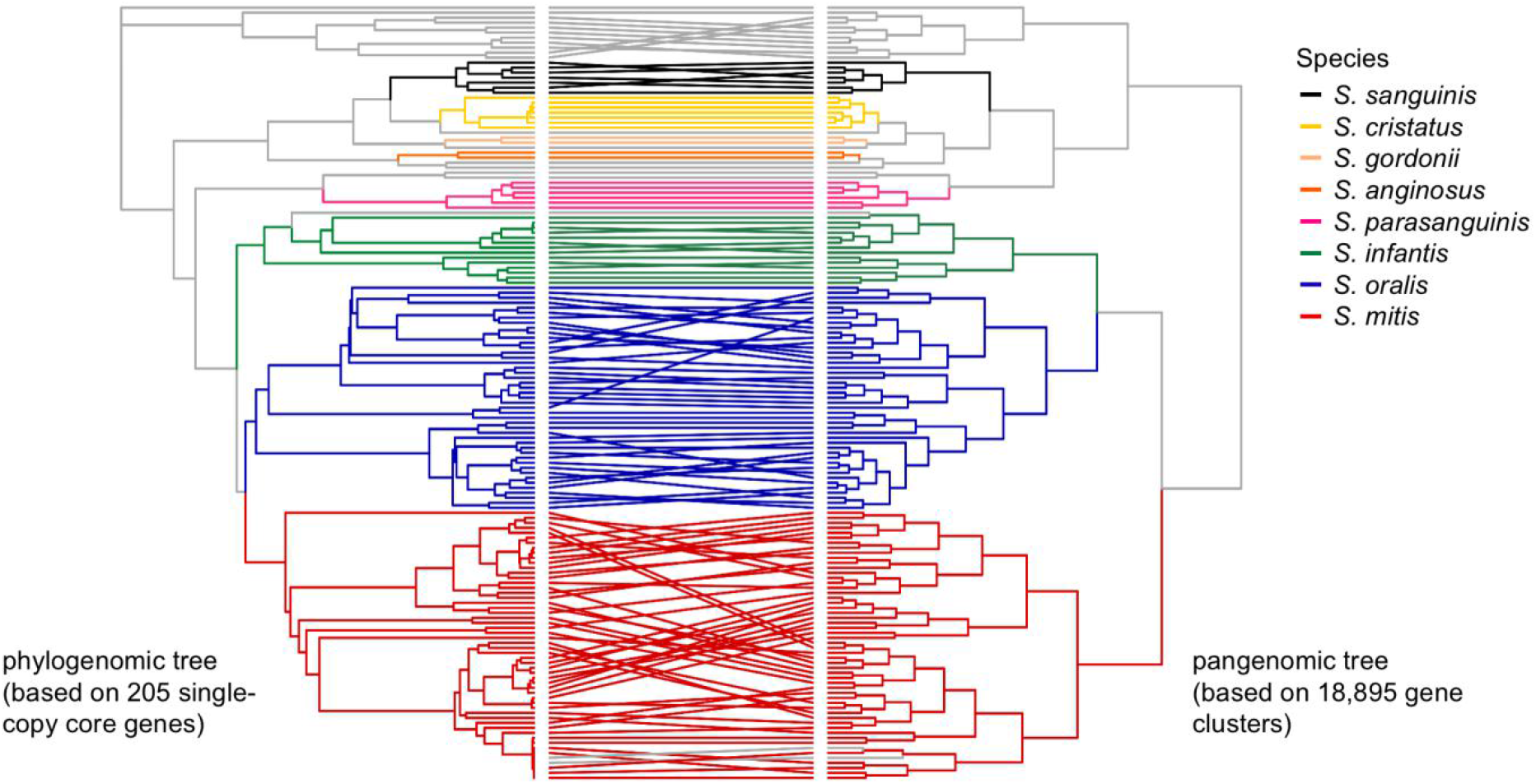
Analysis of phylogeny and analysis of gene content produce congruent results and cluster genomes into the same species-level groups. The phylogenomic tree was constructed using 205 single-copy genes core to the oral streptococci. The pangenomic tree was constructed using the frequencies with which each of the 18,895 genes is present in each genome. Lines connect the end nodes that represent the same genome. Colored boxes indicate species-level clades that contain multiple genomes.

**Figure S4:**
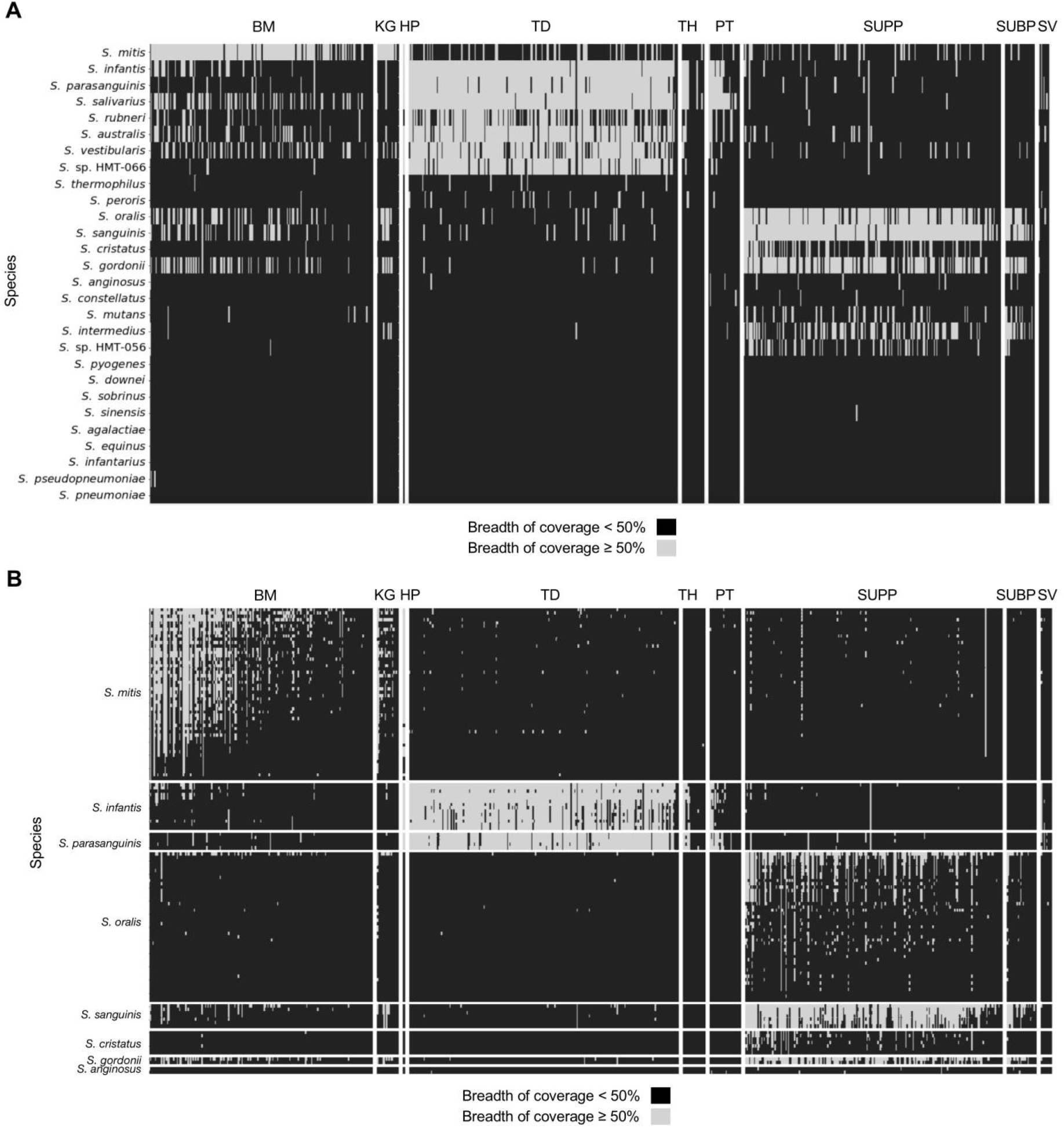
Breadth of coverage also varies for species between oral sites and between strains of the same species within an oral site. The matrices show the breadth of coverage for the oral *Streptococcus* species (A) and individual strains from species with more than one representative genome (B) for each of the metagenomes sampled across nine oral sites. Species or strains with a breadth of coverage of ≥ 50% for a sample are denoted by black and species or strains with a breadth of coverage < 50% are denoted by gray. Species with multiple reference genomes were considered to have a breadth of coverage ≥ 50% if at least one of their representative genomes had a breadth of coverage ≥ 50%. There are 183 buccal mucosa (BM), 23 keratinized gingiva (KG), 1 hard palate (HP), 220 tongue dorsum (TD), 21 throat (TH), 31 palatine tonsils (PT), 209 supragingival plaque (SUPP), 32 subgingival plaque (SUBP), and 8 saliva (SV) samples. The samples are grouped by site and then ranked by descending number of total reads. The strains are first grouped by species and then ranked by descending mean relative abundance across the site (BM, TD, or SUPP) where they are most abundant.

**Figure S5:**
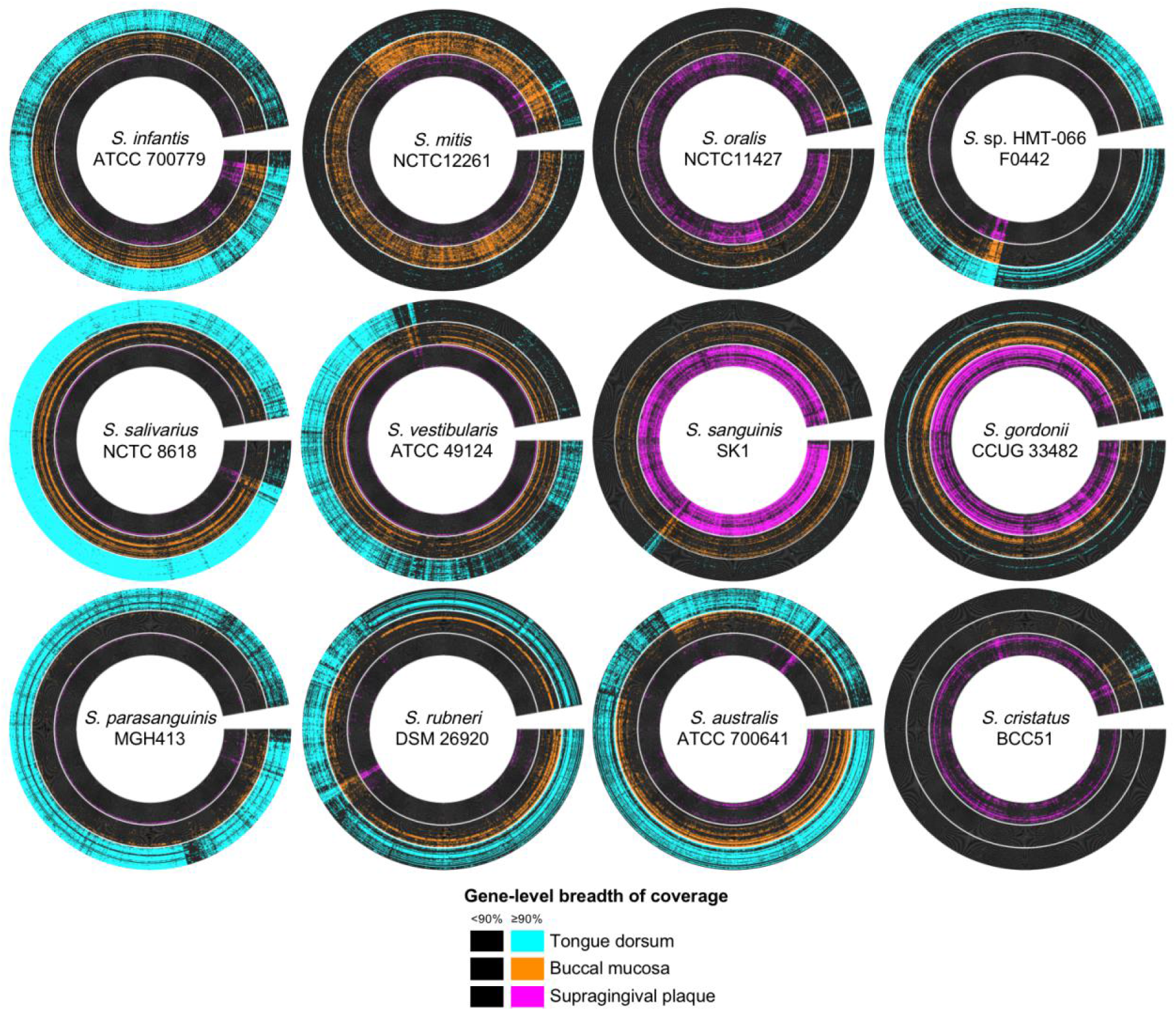
Breadth of coverage validates site tropisms and indicates how well the sequenced genome matches the gene content of the population in the mouth. The radial heatmap displays breadth the of coverage of the predicted open reading frames (ORF) from a representative genome from the 30 buccal mucosa, tongue dorsum, and supragingival plaque samples with the most quality-filtered reads. Each radius represents a predicted ORF. Each concentric ring represents a metagenomic sample. ORFs are black if their breadth of coverage is < 90% and color-coded by site if their breadth of coverage is ≥ 90%. Representative genomes are shown for species with a mean relative abundance ≥ 2% in at least one site. The genomes displayed here represent the genomes from each species which recruited the most reads across all metagenomes and whose NCBI species matched our new species designations.

### *S. pneumoniae* marker sequence

ATGAGTACAAAATATTTATTTATTTACAATGAGATTCGTGAAAAGATTCTTTGTAATAAATATACCATGAACGAACAATTGCCTGATGAAATGACATTAGCTAAACAGTTTGCCTGTAGTCGAATGACGATCAAAAAAGCTTTAGACTTGTTAGTTTCTGAGGGCTTAATTTTTAGAAAACGTGGGCAGGGAACCTTTGTTCTCTCTCGTGGCAGCTCAAAAAGAAAATTAATCGTTCCAGAAAGAGATATCCGGGGACTGACAAAAATATCTGAAGATGCTCATTCTACAATTGACTCGAGGATTATTCACTTCAAATTAGAATTTGCAAATGAATTTTTAGCAGAAAAACTACAGGTCGCTTTGCAGAGTCCAGTTTATAATATTTACCGCCTGCGTATTATTGACGGTAAACCTTATGTTCTGGAACAAACTTATATGAGTACCGATGTTATTCCAGGTA TTACTGAAGATATTTTACAAAAATCGATTTACAATTACATTGAAGGAAAGTTAGGATTGCATATTGCCAGTGCTACAAAAATCTTACGAGCTTCTTCTAGTTCAGAAAATGAGCAACATTACTTGCAGCTCCTTCCAACGGAACCGGTATTTGAAGTAGAACAAGTGGCTTATTTGGATAACGGAACTCCGTTTGAGT ACTCGATTAGTCGTCATCGCTATGATTTATTTGAATTTAATTCTTTTGCATTACGACATTCCTCCTAG

### *S. pseudopneumoniae* marker sequence

ATGTATTACATGAAAAATGAAAATGTTAAGATTTTAATTTGTGAAGATGACTCTTCCGTTAACAGACTTTTATCCTTAGCAATGGAAGTTGAAGGTTATCATTATGTATCAGTTCGGACTGGAGAGGAAGCTTTGCGTCAGATCATTTCGCAATTTCCAGATTTATTATTATTGGATTTGGGTTTGCCAGATATGGATGGTAAAGACATTATTGACAAGATTCGTAGCTTTTCACAGCTACCTGTTATTGTTGTTAGTGCACGTGGAGAAGAAAGTGACAAGATTGATGCACTTGATGCTGGGGCAGATGATTATTTGACGAAACCCTTTAGCATTGATGAGCTTTTCGCTCGGTTAAGAGTTAGTCTTAGGAGGTCAAAGCAGATTAATCAACAAAGTGACGGTAATTCTGAAAAATCATCTTTTACTAATGGCTGGCTACATGTTGATTTTTTATCTAATCGTGTATTTGTTAATAACCAAGAAATTCACTTAACCCCGATTGAGTATAAGTTGCTTTGTCTTCTATCAGAGAATGTTGATAGAGTGTTGACTTATCGTTTTATTGTCAAGGAAATTTGGGGATATTATGAGGAAGATTTTTCTGCTTTGAGAGTTTTTGTTAATACATTGCGAAAAAAAATCGAATTAGGATTGGGTTACTCTAAAATGGTTCAAACTCATATTGGTATCGGTTATCGTATGATTAAGATTGAAAATTATGATGAC AAATAA

## Supplemental Methods

### Mapping Specificity Test

To evaluate the specificity of mapping to the reference genome set, we generated a set of simulated paired-end read samples using the program ‘reads-for-assembly’ (https://github.com/merenlab/reads-for-assembly). Each sample used a single genome from one of the following three categories as a template for the simulated reads — oral streptococci type strain genomes from the reference genome set, oral streptococci genomes not in the reference genome set with ≥ 95% ANI, type strain genomes from other major human oral genera (Eren et al. 2014, Mark Welch et al. 2019). The samples contained 100 bp long reads which had a mean offset of 30 bp with a standard deviation of 1 bp. The reads covered their template genome to a mean depth of 100 reads and had an average base substitution error rate of 0.5%. This error rate falls within the expected range for Illumina reads quality-filtered by low-quality end trimming; the insertion-deletion error rate would be expected to be negligible for these reads (Minoche et al. 2011, Schirmer et al. 2016). We competitively mapped the simulated samples to the dereplicated reference genome set and profiled the samples as described for the real HMP samples.

